# SARS-CoV-2 exposure in wild white-tailed deer (*Odocoileus virginianus*)

**DOI:** 10.1101/2021.07.29.454326

**Authors:** Jeffrey C. Chandler, Sarah N. Bevins, Jeremy W. Ellis, Timothy J. Linder, Rachel M. Tell, Melinda Jenkins-Moore, J. Jeffrey Root, Julianna B. Lenoch, Suelee Robbe-Austerman, Thomas J. DeLiberto, Thomas Gidlewski, Mia K. Torchetti, Susan A. Shriner

**Affiliations:** USDA/APHIS/WS National Wildlife Research Center; Fort Collins, USA; USDA/APHIS/VS National Veterinary Services Laboratories; Ames, USA; USDA/APHIS/WS National Wildlife Disease Program; Fort Collins, USA

## Abstract

Widespread human SARS-CoV-2 infections combined with human-wildlife interactions create the potential for reverse zoonosis from humans to wildlife. We targeted white-tailed deer (*Odocoileus virginianus*) for serosurveillance based on evidence these deer have ACE2 receptors with high affinity for SARS-CoV-2, are permissive to infection, exhibit sustained viral shedding, can transmit to conspecifics, and can be abundant near urban centers. We evaluated 624 pre- and post-pandemic serum samples from wild deer from four U.S. states for SARS-CoV-2 exposure. Antibodies were detected in 152 samples (40%) from 2021 using a surrogate virus neutralization test. A subset of samples was tested using a SARS-CoV-2 virus neutralization test with high concordance between tests. These data suggest white-tailed deer in the populations assessed have been exposed to SARS-CoV-2.

**One-Sentence Summary:** Antibodies to SARS-CoV-2 were detected in 40% of wild white-tailed deer sampled from four U.S. states in 2021.

## Main Text

SARS-CoV-2, the virus that causes COVID-19 in humans, can infect multiple domestic and wild animal species (*1–7*). Thus, the possibility exists for the emergence of new animal reservoirs of SARS-CoV-2, each with unique potential to maintain, disseminate, and drive novel evolution of this virus. Of particular concern are wildlife species that are both abundant and live in close association with human populations (*5*).

The pathogen pressure produced by significant human infections combined with susceptible wildlife hosts at the wildlife-human interface has led multiple authors to signal an urgent call for pro-active wildlife surveillance for early detection of potential reverse zoonosis (spillback) of SARS-CoV-2 into wildlife populations (*8–10*). Reverse zoonosis could lead to the establishment of novel wildlife reservoirs outside of southeast Asia (*11*), a potential that poses significant risks to both human and animal health (*8–10, 12*). Besides health impacts to wildlife, persistent infections in a novel host could lead to adaptation, strain evolution, and re-emergence of strains with altered transmissibility, pathogenicity, and vaccine escape. Cross-species transmission to other wildlife species and the concomitant risks is also a concern (*8, 10*).

*In silico* analyses of SARS-CoV-2 spike protein affinity to the primary host cellular receptor, angiotensin-converting enzyme 2 (ACE2) suggest that multiple animal species endemic to the U.S. are potentially susceptible to SARS-CoV-2 (*13*), including white-tailed deer (*Odocoileus virginianus*). The geographic distribution of this species encompasses most of North America and these animals are particularly abundant near urban population centers located in the eastern U.S. (*14*). Moreover, white-tailed deer can form social groups, a contact structure with the potential to support intraspecies transmission of multiple pathogens (*15*). A SARS-CoV-2 experimental infection study of white-tailed deer showed these deer exhibit sub-clinical infections, shed virus in nasal secretions and feces, and can transmit the virus to contact control deer (*1*). Further, SARS-CoV-2 neutralizing antibodies were detected by seven days post-inoculation (dpi) and were consistently observed through 21 dpi (*1*).

Surveillance prioritization for early detection of potential SARS-CoV-2 reverse zoonosis should be based on risk assessment that considers ACE2 heterogeneity, potential for human interaction, infection dynamics, probability of onward transmission, behavior, and contact networks (*9, 10*). As reviewed elsewhere (*10*), cervids are a high priority across each of these characteristics.

The USDA/APHIS/Wildlife Services National Wildlife Disease Program (NWDP) conducts wildlife disease surveillance for a variety of pathogens and species throughout the U.S. In January 2021 we leveraged this resource to initiate a pilot serosurveillance program for SARS-CoV-2 exposure in white-tailed deer. While serological testing primarily detects historical infection, the extended period for detecting antibodies, compared to specific viral or molecular detection of the pathogen, significantly increases the probability of detection due to the prolonged duration of circulating antibodies (*10, 16*). In the case of emerging pathogens such as SARS-CoV-2, serosurveys also demonstrate absence of exposure to provide baseline information for surveyed populations prior to pathogen emergence. Here, serum samples were collected opportunistically as part of wildlife management activities (e.g., surveillance for chronic wasting disease and bovine tuberculosis, and urban removals) as a preliminary survey to evaluate the potential role of free-ranging white-tailed deer in the epidemiology of SARS-CoV-2.

From January-March 2021, we received 385 wild white-tailed deer serum samples from four states: Michigan (MI, N = 113), Pennsylvania (PA, N = 142), Illinois (IL, N = 101), and New York (NY, N = 29, Table 1). The NWDP maintains extensive wildlife sample archives across multiple species, including serum samples collected for cervid surveillance. We used this repository to select 239 wild white-tailed deer serum samples from 2011-2020 (pre- and early pandemic) from five states for serological analyses: IL (N = 16), MI (N = 37), PA (N = 104), New Jersey (N = 8), and NY (N = 74). To the extent possible, archive samples were matched to samples collected in 2021 with respect to state and county to serve as controls to identify potential endemic coronaviruses that might cross-react in laboratory testing. The majority of archive samples were from 2018-2020 (N = 182).

**Table 1.** Test concordance for the Genscript cPass™ surrogate virus neutralization test (sVNT) and the virus neutralization test (VNT).

|  | VNT<br>Positive | VNT<br>Negative |
| --- | --- | --- |
| sVNT<br>Positive | 24 | -- |
| sVNT<br>Negative | -- | 24 |

All samples were screened at the National Wildlife Research Center (Fort Collins, USA) using a surrogate virus neutralization test (sVNT, Genscript cPass™). This test is species independent and allows for testing in biosafety level 2 laboratories, making it an appropriate choice for high throughput screening of wildlife samples. The sVNT detects total neutralizing antibodies (as measured by percent inhibition) that interfere with the affinity of the SARS-CoV-2 spike protein receptor binding domain to ACE2 (*17*). Because the sVNT has not been validated for deer, we also conducted parallel testing at the National Veterinary Services Laboratories (Ames, USA) on a subset of samples using a highly specific SARS-CoV-2 virus neutralization test (VNT) run with infectious SARS-CoV-2.

Antibodies to SARS-CoV-2 were detected in 40% of the 2021 surveillance samples (Table 1). Antibodies were also detected in three samples from 2020 and one sample from 2019. No detections were observed in samples from 2011-2018. The results from the sVNT screening showed high concordance with those obtained by VNT (Table 1). Specifically, 24/24 of 2021 detections and 24/24 of 2021 negatives were concordant for sVNT compared to VNT.

Most of the positive samples from 2021 had percent inhibition values between 80-100% while the 2019-2020 positive samples had relatively low percent inhibition values (30.03-43.72, Figure 1). Percent inhibition scores ≥30 are considered positive for this assay. Low percent inhibition could represent potential waxing/waning immunity, non-specific antibody binding, or cross-reactivity from exposure to unknown endemic coronaviruses. The three positive samples from 2020 were collected in January, very early in the pandemic. In fact, the majority of the 2020 samples that were available for testing were from January-March, with only 21 samples collected later in the year, 20 of which were collected in October from a single location. Consequently, we have limited information on prevalence over time in 2020.

**Fig. 1.**
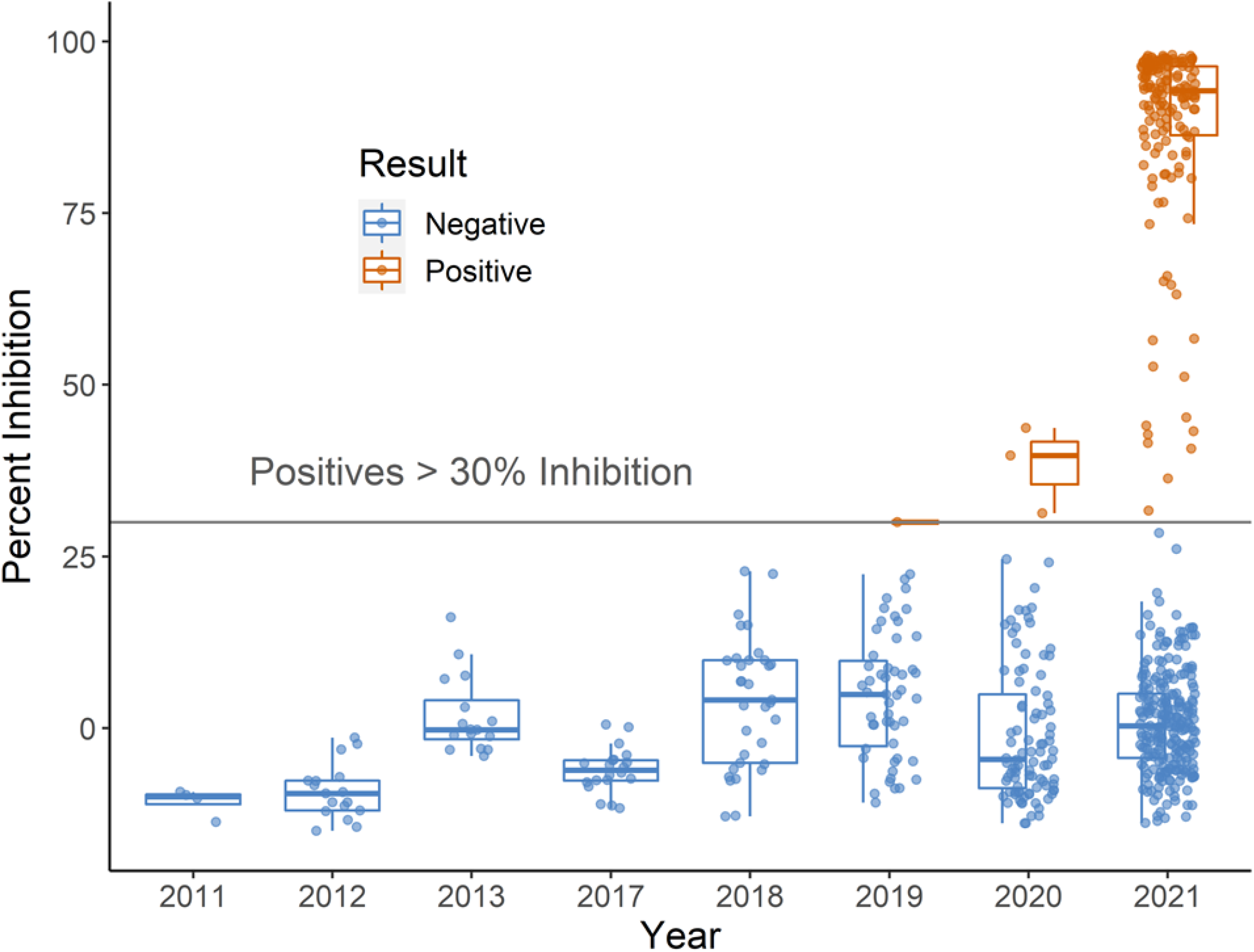
SARS-CoV-2 serological results for white-tailed deer. Serum samples were tested using the Genscript cPass™ surrogate virus neutralization test (sVNT).

Seroprevalence in free-ranging white-tailed deer varied by county and state (Table 2, Figure 2). Considering only 2021 samples, at the state level, the lowest seroprevalence observed was 7% in Illinois and the highest was 67% in Michigan with intermediate seroprevalence in New York (31%) and Pennsylvania (44%). Seroprevalence for individual counties was highly clustered with nearly half of the 32 counties sampled showing no evidence of SARS-CoV-2 exposure.

**Table 2.** County level seroprevalence for SARS-CoV-2 in white-tailed deer sampled January-March, 2021.

| State | County | N | #Pos | Sero-<br>prevalence | State | County | N | #Pos | Sero-<br>prevalence |
| --- | --- | --- | --- | --- | --- | --- | --- | --- | --- |
| MI | Emmett | 3 | 0 | 0 | IL | Dekalb | 1 | 0 | 0 |
| MI | Lenawee | 5 | 0 | 0 | IL | Ogle | 2 | 0 | 0 |
| MI | Montmorency | 7 | 0 | 0 | IL | Kane | 5 | 0 | 0 |
| MI | Jackson | 5 | 2 | 40 | IL | Will | 11 | 0 | 0 |
| MI | Presque Isle | 12 | 6 | 50 | IL | Winnebago | 11 | 0 | 0 |
| MI | Alpena | 34 | 25 | 74 | IL | LaSalle | 15 | 0 | 0 |
| MI | Alcona | 21 | 18 | 86 | IL | Kendall | 18 | 0 | 0 |
| MI | Mecosta | 11 | 10 | 91 | IL | Grundy | 19 | 0 | 0 |
| MI | Gratiot | 5 | 5 | 100 | IL | Kankakee | 6 | 1 | 17 |
| MI | Ingham | 5 | 5 | 100 | IL | Cook | 9 | 3 | 33 |
| MI | Isabella | 5 | 5 | 100 | IL | McHenry | 3 | 2 | 67 |
| PA | Wayne | 11 | 0 | 0 | IL | Livingston | 1 | 1 | 100 |
| PA | Warren | 14 | 0 | 0 | NY | Suffolk | 8 | 0 | 0 |
| PA | Westmoreland | 14 | 0 | 0 | NY | Onondaga | 21 | 9 | 43 |
| PA | Montgomery | 54 | 22 | 41 |  |  |  |  |  |
| PA | Philadelphia | 16 | 7 | 44 |  |  |  |  |  |
| PA | Huntingdon | 5 | 5 | 100 |  |  |  |  |  |
| PA | Snyder | 28 | 28 | 100 |  |  |  |  |  |
| **OVERALL** |  |  |  |  |  |  | **385** | **154** | **40** |

**Fig. 2.**
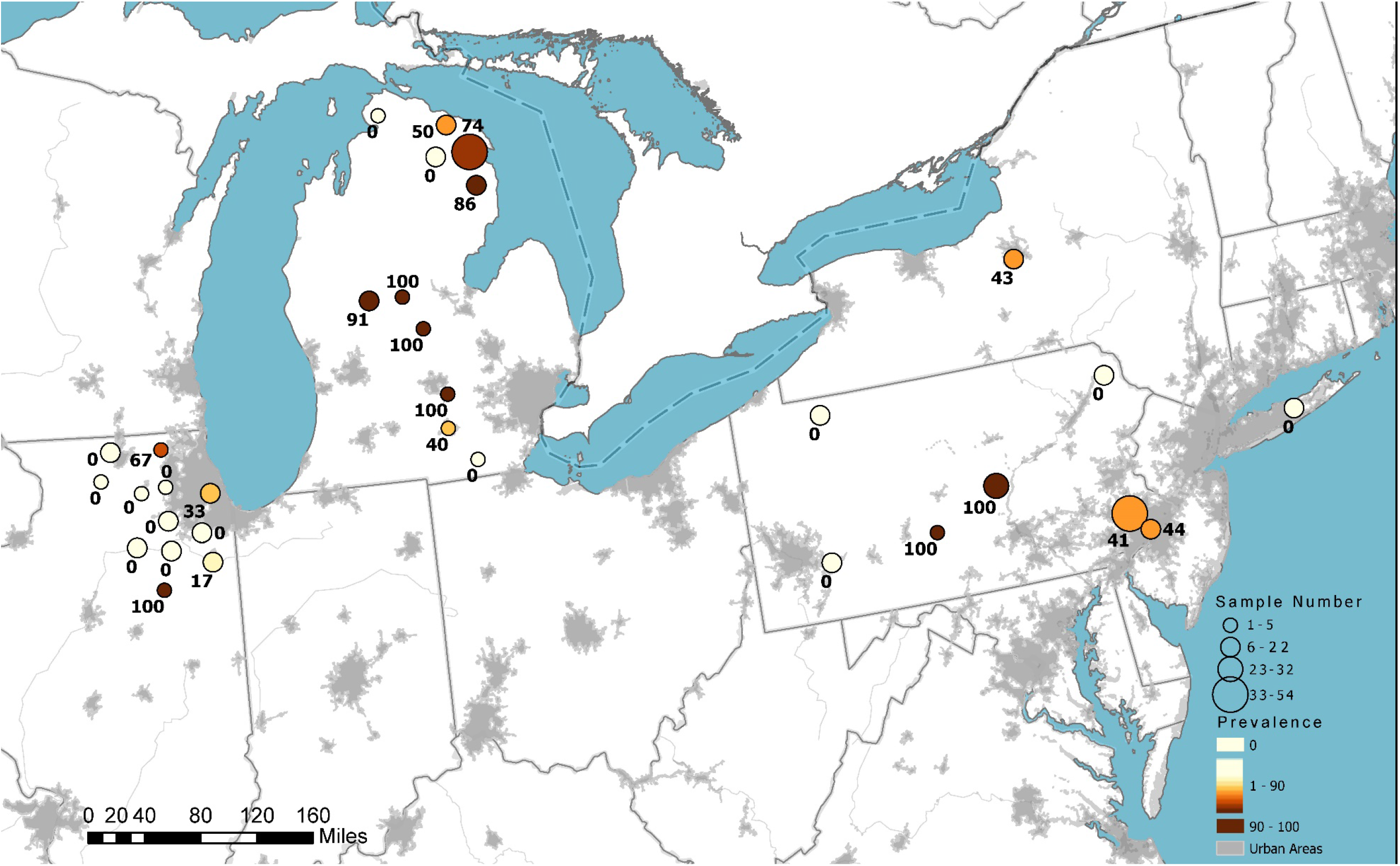
SARS-CoV-2 antibody detection in white-tailed deer sampled in 2021 in the United States. Circle size indicates the relative number of samples tested. Color represents the relative seroprevalence, and numbers are the specific prevalence in a county.

An important consideration is the potential for assay cross-reactivity with antibodies against other coronaviruses. In analyses of human serum samples, the sVNT achieved 99.93% specificity and 95-100% sensitivity (*17*). No cross-reactivity was observed for several human coronaviruses, including 228/NL63, OC43, or MERS. Minor cross-reactivity was detected between SARS and SARS-CoV-2 (*17–19*). In contrast, no cross-reactivity has been identified in SARS-CoV-2 specific VNT for closely related human coronaviruses (*17–19*) or animal viruses (*20*).

Limited work has been done on coronaviruses in white-tailed deer for baseline information on potential cross-reactivity. Previous studies have identified bovine-like coronaviruses in cervids in the U.S. (*21, 22*). However, differences in receptor affinity of these viruses, genetic variability, and previous evaluations of serological cross-reactivity in coronaviruses suggest limited potential for cross-reactivity to antibodies to the SARS-CoV-2 spike protein receptor binding domain (*21–23*).

Several potential transmission routes are possible for movement of this virus into wild deer populations. In the case of SARS-CoV-2 outbreaks in farmed mink, direct transmission of the virus from infected humans to mink is the only definitive transmission route identified to-date (*24, 25*). Multiple activities could bring deer into contact with people, including captive cervid operations, field research, conservation work, wildlife tourism, wildlife rehabilitation, supplemental feeding, and hunting (*10*). Wildlife contact with contaminated water sources has also been offered as a potential transmission route (*11*), although transmissibility of SARS-CoV-2 from wastewater has yet to be conclusively demonstrated (*26*). Transmission from fomites or other infected animal species cannot be discounted.

Overall, these results indicate that the white-tailed deer populations examined in MI, PA, IL, and NY were exposed to SARS-CoV-2. While neutralizing titers for some of the samples were relatively high and suggestive of infection (Supp. Info), we cannot confirm infection based on serological data. These results emphasize the need for continued and expanded wildlife surveillance to determine the significance of SARS-CoV-2 in free-ranging deer. We also recommend SARS-CoV-2 surveillance of susceptible predators and scavengers that have a high probability of interacting with deer. Future wildlife surveillance should incorporate methods specifically designed to detect, isolate, and genetically characterize SARS-CoV-2 and to identify potential variants, as well as other endemic coronaviruses. These methods, combined with dedicated wildlife disease surveillance programs, can shed light on how zoonotic pathogen spillback into novel wildlife reservoirs may affect pathogen adaptation, evolution, and transmission.

## Acknowledgments

We thank the USDA Wildlife Services biologists and field specialists who coordinated and collected samples that made this work possible: **Dustin Arsnoe, Mark Jackling, Mitch Oswald, Kyle Van Why**, Brody Allen, Dakota Bird, Caleb Brown, Charles Cini, Craig Hicks, J. Nave, Rex Schanck, Trent Speaks. The findings and conclusions in this publication are those of the authors and should not be construed to represent any official U.S. Government determination or policy.

## Author contributions

Conceptualization: SAS

Methodology: TJL, SNB, JJR, MKT, SAS

Investigation: JCC, RMT, MJM, JWE, MKT, SAS

Project administration: SR-A, TJD, JBL, TG, MKT, SAS

Writing – original draft: JCC, SAS, SNB, MKT

Writing – review & editing: JCC, RMT, TJL, JWE, SNB, JJR, SR-A, TJD, JBL, TG, MKT, SAS

## Data and materials availability

All data files are available from the https://www.fs.usda.gov/rds/archive/database.

## Supplementary Materials

### Materials and Methods

The USDA/APHIS/Wildlife Services National Wildlife Disease Program (NWDP) conducts wildlife disease surveillance for a variety of pathogens and species throughout the U.S. We leveraged this resource to initiate a sero-surveillance program for SARS-CoV-2 exposure in white-tailed deer in January 2021. Serum samples were collected opportunistically as part of operational activities conducted by NWDP biologists (e.g., surveillance for chronic wasting disease and bovine tuberculosis, urban removals). All samples were shipped to the USDA National Wildlife Research Center (Fort Collins, CO USA) where they were received, logged, and placed in a −80°C ultracold freezer until testing. Samples were tested using the species and isotype independent Genscript cPass™ surrogate virus neutralization test (sVNT) in an ELISA format. The assay was conducted according to manufacturer’s instructions.

In addition to the samples collected specifically for SARS-CoV-2 screening, we also tested archived white-tailed deer serum samples from Wildlife Services biological archive using the sVNT. In general, archive samples were selected from the same state and counties as the 2021 SARS-CoV-2 surveillance samples except for a small portion of samples that were selected from nearby counties (as available) as well as ten samples from New Hampshire and New Jersey.

A subset of samples was submitted to the USDA National Veterinary Services Laboratories (NVSL, Ames, IA USA) for testing by a SARS-CoV-2 specific virus neutralization test. The subset included 48 surveillance samples from 2021 (24 positives and 24 negatives by sVNT).

